# Identification of hemicatenane-specific binding proteins by fractionation of Hela nuclei extracts

**DOI:** 10.1101/844126

**Authors:** Oumayma Rouis, Cédric Broussard, François Guillonneau, Jean-Baptiste Boulé, Emmanuelle Delagoutte

## Abstract

DNA hemicatenanes (HCs) are DNA structures in which one strand of a double stranded helix passes through the two strands of another double stranded DNA. Frequently mentioned as DNA replication, recombination and repair intermediates, they have been proposed to participate in the spatial organization of chromosomes and in the regulation of gene expression. To explore potential roles of HCs in genome metabolism, proteins capable of binding specifically HCs were purified by fractionating nuclear extracts from Hela cells. This approach identified three RNA-binding proteins: the Tudor-Staphylococcal Nuclease Domain 1 (SND1) protein and two proteins from the Drosophila behavior human splicing family, the ParaSpeckle Protein Component 1 and the Splicing Factor Proline- and Glutamine- rich protein. Since these proteins were partially pure after fractionation, truncated forms of these proteins were expressed in *E. coli* and purified to near homogeneity. The specificity of their interaction with HCs was re-examined *in vitro*. The two truncated purified SND1 proteins exhibited a high specificity for HCs, suggesting a role of SND1 protein in targeting the basic transcription machinery to HC structures.

## Introduction

DNA hemicatenane (HC) consists of a junction of four DNA strands in which one strand of one double helix passes in between the two strands of another double helix (Fig 1A). They can implicate one or two different double stranded DNA molecules and, in this later case, the two DNA helices may or may not be homologous. DNA HCs might appear at several times during the cell cycle. For instance, the convergent migration of two Holliday junctions formed during DNA recombination may lead to a HC depending on the relative topology of the two junctions. HC may be created during the initiation [1] and termination [2] phases of DNA replication or when replication is inhibited [3]. Their presence close to a replication fork in progress may help repair and restart a stalled DNA replication fork, due to the proximity of the newly synthesized daughter DNA molecules [1]. Finally, the ultra fine bridges that can be observed during the anaphase step of mitosis after a perturbed or incomplete DNA replication may consist of two homologous chromosomes topologically linked by HCs (for recent reviews see [4–6]).

**Figure 1.**
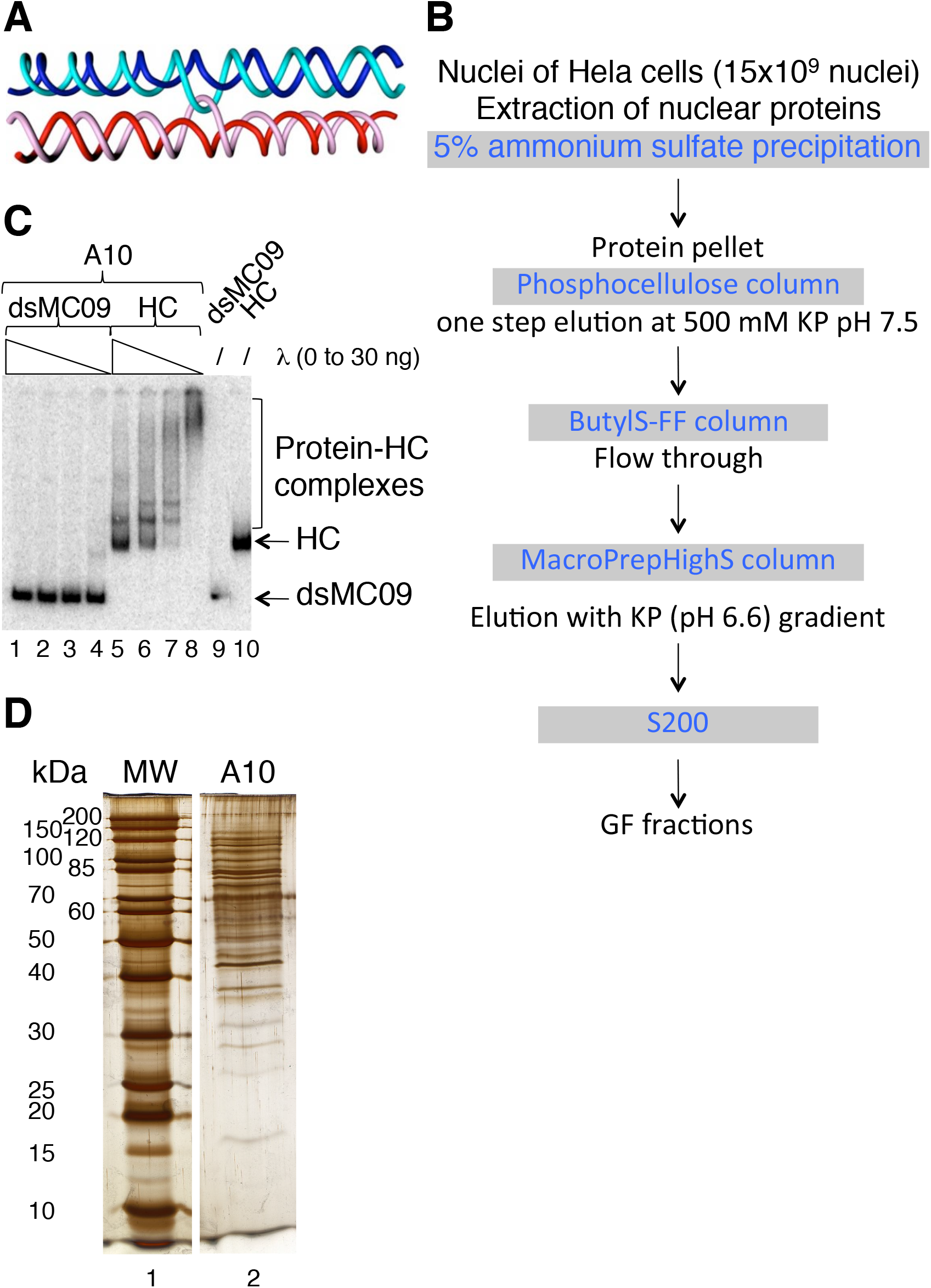
Fractionation of Hela nuclear extracts. A Schematic representation of a DNA hemicatenane. In this scheme, the two double stranded DNAs that are topologically linked by a hemicatenane are not homologous and therefore are represented by two different colors: blue and red. The light and dark blue strands are complementary, as the light and dark red strands. B Scheme of the procedure of fractionation of Hela nuclear extracts. The procedure starts with an ammonium sulfate precipitation that is followed by four chromatographies on different media, as indicated. C Proteins contained in 1 μL of fraction A10 of the size exclusion chromatography were incubated with radiolabeled dsMC09 (lanes 1–4, 9) or HC (lanes 5–8, 10) in the presence of increasing amount of λ DNA (0 (lanes 4 and 8), 5 (lanes 3 and 7), 10 (lanes 2 and 6), 30 (lanes 1 and 5) ng). After incubation, species were resolved by electrophoresis on a native polyacrylamide gel. Free DNA (dsMC09 or HC) and Protein-HC complexes are indicated. D Analysis of the fraction A10 of the size exclusion chromatography by electrophoresis on a polyacrylamide gel followed by a silver staining of the gel. Lane 1: MW; the size of the proteins is given in kDa; lane2: 30 μL of fraction A10.

Faithful and accurate processing of HCs is essential for genome stability. For HCs resulting from the convergent migration of two Holliday junctions, the process of unlinking the two DNA molecules of the HC is called dissolution (for reviews see [7,8]). In humans, the dissolvasome complex (BTRR) is responsible for the branch migration and dissolution reactions. In this complex, the RecQ-like Bloom helicase (B in BTRR) uses its ATPase activity to migrate the Holliday junctions [9,10] and its oligomeric structure may be responsible for the convergent migration of two Holliday junctions [7]. The Topoisomerase IIIα (T in BTRR), a type IA topoisomerase, through a transesterification and strand passage reaction, decatenates the HC. Its activity is stimulated by the RecQ-mediated instability, RMI1 and RMI2 (RR in BTRR), factors [11–13]. Devoid of catalytic activity, the RMI1 might stabilize the open form of the Topoisomerase IIIα to favor strand passage. Finally, one family of ultra fine bridges has been shown to be decorated by the BTRR complex and it is possible that the decatenase activity of Topoisomerase IIIα be used to resolve their inter-strand links [14,15].

*In vitro* experiments have shown that re-association of DNA fragments containing the polyCA/polyTG repeat sequences leads to the formation of an alternative structure that has been proposed to be a HC [16]. The High Mobility Group Box 1 and 2 (HMGB1 and HMGB2) proteins stimulate this process [17,18]. Following this observation, it has been proposed that HCs may participate in the genome organization and in the regulation of its transcription [19]. In this model, genome is organized in large loops maintained at their base by topological knots, such as HCs. Unless stabilized, knots can migrate and their location on the chromosome can change. Through the possibility of knots migration and stabilization, chromosomal loops may expose a variety of different DNA sequences, making thus possible the expression of different genes. Inherent to this model is the reversible and transient stabilization of the knots, either by a specific sequence or by some HC-specific proteins.

Our goal in this work was to explore further these hypotheses by seeking nuclear proteins capable of stabilizing HC structures by specifically binding to them. We took advantage of the existence of a procedure to build HCs from two distinct double stranded DNA mini-circles (dsMCs) [20] to fractionate nuclear extracts from Hela cells over several chromatographic media. This approach made possible the identification of three RNA-binding proteins capable of binding specifically to HC: the Tudor-Staphylococcal Nuclease Domain 1 (SND1) protein and two proteins from the Drosophila behavior human splicing (DBHS) family, the ParaSpeckle Protein Component 1 (PSPC1) and the Splicing Factor Proline- and Glutamine- rich protein (SFPQ). Since these proteins were partially pure after fractionation of the Hela nuclear extracts, and because of the difficulty to purify them in a full length form, truncated forms of these proteins were expressed in *E. coli* and purified to near homogeneity. Specificity of the interaction between the purified proteins and HCs was re-examined in a gel shift assay. The results of our characterization suggest a possible role of SND1 protein as a transcription factor targeting the basic transcription machinery to HC structures.

## Results and discussion

Our search for HC-specific binding proteins required to have available radiolabeled DNA HCs. These DNA structures were built as described in [20]. The starting material was two distinct radiolabeled dsMCs, one of 215 base pairs (dsMC10) and one of 235 base pairs (dsMC09), produced after circularization of ^32^P-end labeled fragments (Fig EV1). The nicking of the dsMCs at specific sites provided linear single stranded fragments of 215 (L10ss) and 235 (L09ss) nucleotides, and circular single stranded DNAs of 215 (C10ss) and 235 (C09ss) nucleotides. The circularization of the linear single stranded L09ss and L10ss fragments under conditions that kept them in close proximity led in part to single stranded catenanes. C09ss and C10ss were re-annealed on the single stranded catenanes with the wheat germ topoisomerase I to finally give DNA HCs (Fig EV1).

### Fractionation of Hela nuclei protein extracts

To isolate proteins that bind specifically HCs, we first prepared a protein nuclear extract from Hela nuclei cells at 0.6 M NaCl. At this salt concentration, the nuclei chromatin swells and the proteins that fall off the DNA can be recovered by a high-speed centrifugation. We next fractionated this nuclear protein extract following a procedure (Fig 1B) that consisted in five steps: a 5 % ammonium sulfate precipitation followed by four chromatographies: 1) phosphocellulose chromatography, 2) hydrophobic chromatography, cationic exchange chromatography and finally 4) a size exclusion chromatography. Along the fractionation procedure, the fractions were selected for containing HC-specific binding proteins based on a simple and robust binding assay (Fig 1C): for example proteins contained in the fraction A10 of the size exclusion chromatography did bind HC (lanes 5–8 of Fig 1C) but not dsMC09 (lanes 1–4 of Fig 1C). Furthermore, the proteins assembled on HC efficiently resisted the competition with λ DNA since excess of λ DNA did not completely abolish the protein-DNA interaction (lanes 5–8 of Fig 1C). At the end of this 5-step procedure the fractions were not homogeneous (Fig 1D) but greatly enriched in HC-specific binding proteins.

### Identification of HC-specific binding proteins by mass spectrometry

To identify the proteins assembled on the HC, a large scale (150 μL) interaction between HC and the fraction A10 of the size exclusion chromatography was performed, species were resolved by electrophoresis on a polyacrylamide gel and the material that migrated slower than the naked HC was analyzed for its protein content by mass spectrometry (Table 1). Note that (i) two concentrations of competitor λ DNA were tested: 12 or 23 ng (left panel of Fig 2A) and (ii) the material above the naked HC and used for mass spectrometry analysis was divided into two gel pieces (green rectangles on right panel of Fig 2A). Among the proteins identified specifically in the samples containing HC and protein fraction with a good score (Mascot score > 25 and number of peptides ≥ 2) were the ParaSpeckle Protein Component 1 (PSPC1), the Splicing Factor Proline- and Glutamine- rich protein (SFPQ), the Tudor-Staphylococcal Nuclease-like protein (SND1) and the Annexin A2 protein score (Table 1, lanes 8 and 10 of Fig 2A). We analyzed the fraction A11 of the size exclusion chromatography in the same manner (Fig 2B) and found with a good score the PSPC1 and the Polypyrimidine Tract Binding Protein (PTBP1) proteins specifically in the sample containing HC and proteins (Table 1, lane 6 of Fig 2B).

**Table 1.** Names and parameters of identification of HC-binding proteins by mass spectrometry.

| Name of the protein | Name of the gel piece | Mascot score | Peptide number | % of coverage |
| --- | --- | --- | --- | --- |
| Annexin A2 | 10p2-β | 64 | 2 | 5 |
| PSPC1 (ParaSpeckle Protein Component 1) | 10p2-γ | 78 | 3 | 5.9 |
| | 10p1-β | 74 | 3 | 5 |
| | 11p-β | 91 | 2 | 3.6 |
| PTBP1 (Polypyrimidine Tract-Binding Protein 1) | 11p-β | 77 | 3 | 5.1 |
| SFPQ (Splicing Factor Proline Glutamine-rich) | 10p1-β | 97 | 3 | 4.8 |
| SND1 (Staphylococcal Nuclease Domain-like protein 1) | 10p1-β | 98 | 2 | 2.6 |

**Figure 2.**
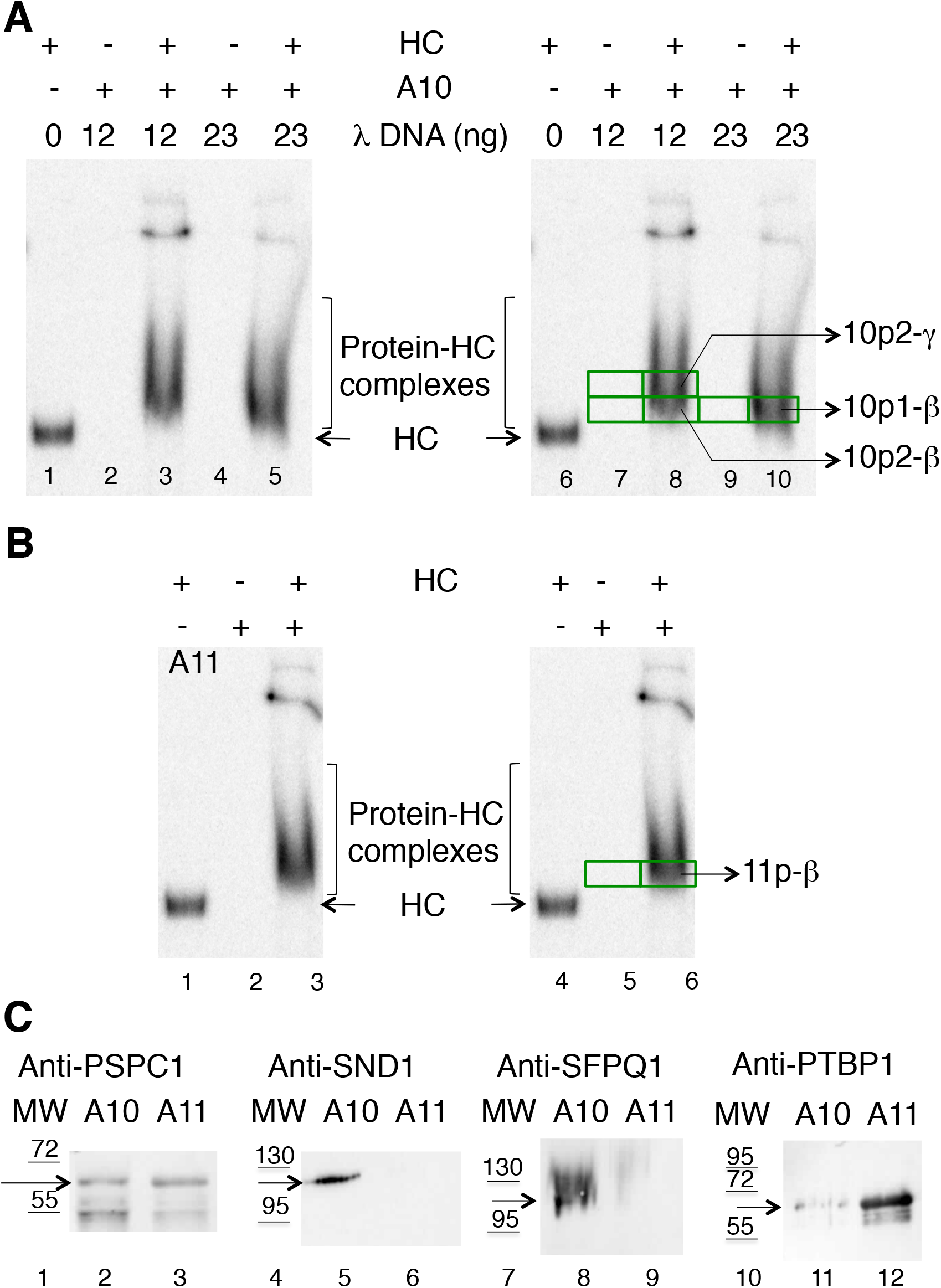
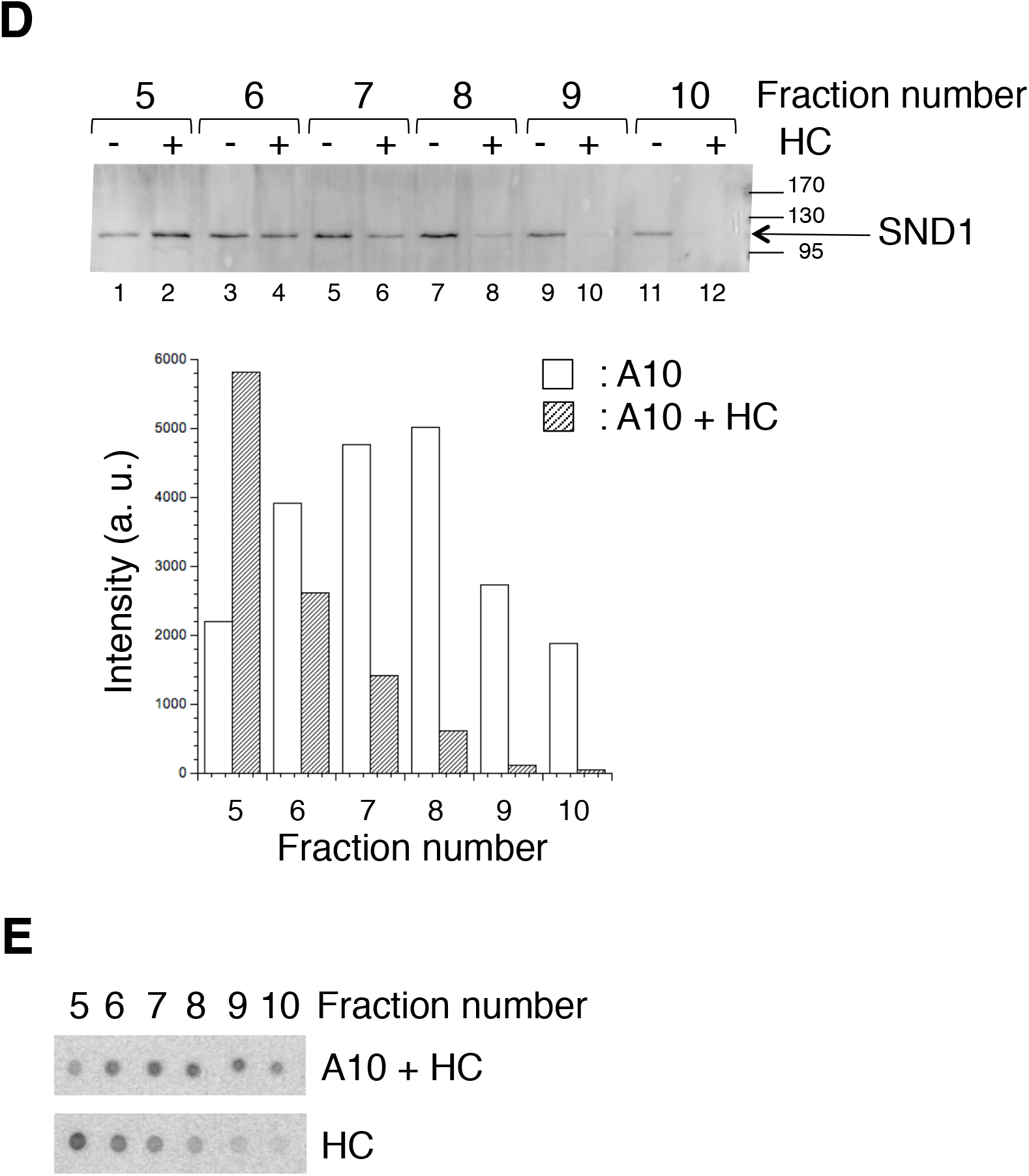
Identification of HC-specific binding proteins in the fractions A10 and A11 of the size exclusion chromatography. A Products of the interaction between radiolabeled HC (16 femtomoles) and proteins of the fraction A10 of the size exclusion chromatography (45 μL) were resolved by electrophoresis on a polyacrylamide gel (lanes 3, 5, 8, 10). 12 (lanes 2, 3, 7, 8) or 23 (lanes 4, 5, 9, 10) ng of λ DNA were added to reduce non specific binding. Sample without protein (lanes 1 and 6) or without HC (lanes 2, 4, 7, 9) were also loaded to serve as control for mass spectrometry analysis. Free DNA (dsMC or HC) and Protein-HC complexes are indicated. On the right panel, the gel pieces that were analyzed by mass spectrometry are shown as green rectangles and the name of the gel pieces of interest is indicated on the right side of the figure. B Same as in (A) but with fraction A11 of the size exclusion chromatography. No λ DNA was added in the reaction mixes. Lanes 1 and 4: HC without fraction A11; lanes 2 and 5: fraction A11 without HC; lanes 3 and 6: HC + fraction A11. On the right panel, the green rectangles represent the two gel pieces that were analyzed by mass spectrometry. The name of the gel piece of interest is indicated on the right side of the figure. C Fractions A10 and A11 were tested for their content of PSPC1, SND1, SFPQ and PTBP1 proteins by western blot. Lanes 1, 4, 7, 10: MW; the size of the proteins is given in kDa; lanes 2, 5, 8, 11: fraction A10; lanes 3, 6, 9, 12: fraction A11. The identity of the antibodies used to probe the faction is indicated above each panel. The arrow points to the protein of interest. D and E Three samples were prepared and centrifuged on a sucrose gradient: sample with HC, sample with fraction A10 and sample with fraction A10 mixed with HC. After centrifugation, 200 μL fractions were collected from the top to the bottom of the gradient. Number of the fraction increases from top to bottom. In (D), fractions 5 to 10 of the sucrose gradients were tested for their SND1 content by western blot (band indicated by an arrow on the right side of the panel). Lanes 1, 3, 5, 7, 9, 11: the fraction A10 was loaded on the sucrose gradient. Lanes 2, 4, 6, 8, 10, 12: the mixture (fraction A10 + HC) was loaded on the sucrose gradient. Bands on the membrane were quantified using Image J software and the relative intensity of each band is plotted as a function of fraction. The white bars correspond to the fraction A10 sample and the hatched bars to the (fraction A10 + HC) sample. In (E), fractions 5 to 10 of the sucrose gradients were tested for their radioactivity content. An aliquot of each fraction (0.5 μL) was spotted on a nitrocellulose membrane. When dried, the membrane was exposed on a ^32^P-sensitive screen. After exposure, the screen was scanned. The radioactivity profile of the HC sample is compared with that of the (fraction A10 + HC) sample.

We focused our interest on three proteins, SND1, PSPC1 and SFPQ for two reasons. Firstly, HCs may participate in the regulation of the genome transcription [19], and secondly, SND1, PSPC1 and SFPQ are key players of the RNA metabolism, including DNA transcription. We checked by western blot for their presence in the size exclusion fractions A10 and A11 (Fig 2C). Results indicated that the three proteins could indeed be detected into the protein fraction A10. Together with the PSPC1 protein, PTPB1 identified in the 11p-β gel piece (Table 1, Fig 2B) could also convincingly be detected in the fraction A11 (Fig 2C). We note that the anti-PSPC1 antibody highlighted two bands in the A10 and A11 fractions, the band of ≈ 65 kDa (indicated by an arrow on lanes 2 and 3 of Fig 2C) corresponding to the full length protein (predicted MW = 59 kDa).

### Characterization of the PSPC1-HC and (PSPC1-SFPQ)-HC nucleoprotein complexes with purified proteins

PSPC1 and SFPQ are two members of the Drosophila behavior human splicing (DBHS) family that includes nuclear proteins implicated in various nuclear functions, such as RNA biogenesis and transport, paraspeckle formation or DNA repair (for a review on DBHS proteins, see [21]). Devoid of catalytic activities but capable of binding a variety of proteins and nucleic acids, members of the DBHS family have been proposed to serve as a “multipurpose molecular scaffold”. DBHS proteins carry a highly conserved DBHS region that consists of two tandem RNA recognition motifs (RRM1 and RRM2 on Fig 3A), a NonA/paraspeckle domain (NOPS on Fig 3A) and a coiled-coil domain. These four domains are responsible for homo- and/or hetero-dimerization of the proteins. They are flanked at their carboxy terminal side by an intrinsically disordered region, G-rich in case of SFPQ protein and GP-rich in case of PSPC1 and a nuclear localization signal (NLS on Fig 3A). The sequences that flank the DBHS core at the amino terminal side of the core are also of low complexity and unstructured. The additional amino terminal domain of PSPC1 is AP-rich whereas that of SFPQ is longer, carries a GPQ-rich domain and a DNA binding region (DBD on Fig 3A) that might function in binding to gene promoters [22] and to DNA damages [23].

**Figure 3.**
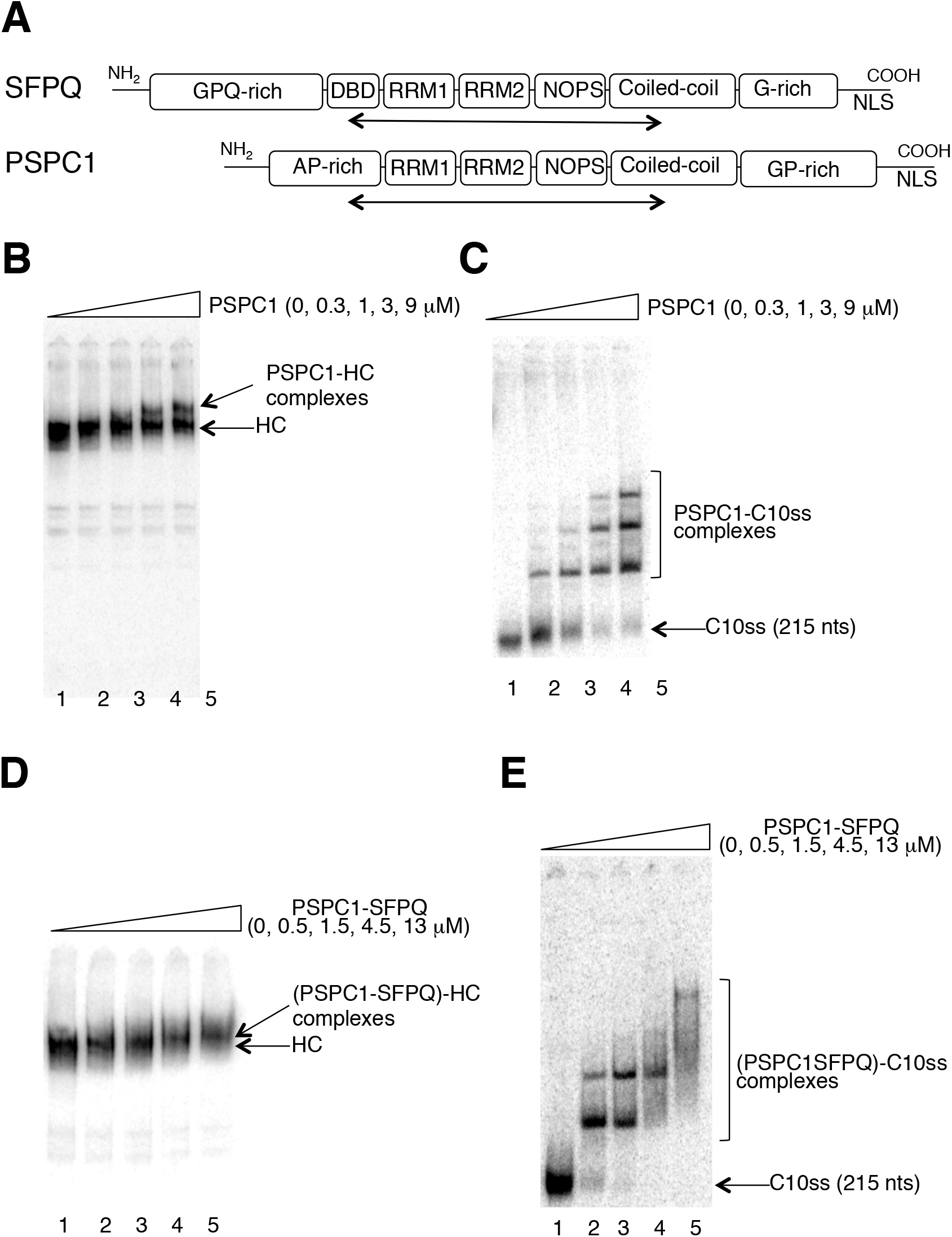
Interaction between purified PSPC1 or PSPC1-SFPQ with DNA. A The organization of SFPQ and PSPC1 in domains is shown. The double arrow underneath each schematic representation corresponds to the truncated protein expressed in *E. coli* and purified. B to E Molecular interactions were performed in a final volume of 7.75 μL with 0.1 femtomole of radiolabeled DNA and the indicated amount of purified protein. Species were resolved by electrophoresis on a polyacrylamide native gel. Free DNAs and bound DNAs are indicated. B The DNA used is HC and the tested protein is PSPC1 homo-dimer. Lane 1: no PSPC1; lane 2: 300 nM PSPC1; lane 3: 1 μM PSPC1; lane 4: 3 μM PSPC1; lane 5: 9 μM PSPC1. C The DNA used is C10ss and the tested protein is PSPC1. Lane 1: no PSPC1; lane 2: 300 nM PSPC1; lane 3: 1 μM PSPC1; lane 4: 3 μM PSPC1; lane 5: 9 μM PSPC1. D The DNA used is HC and the tested protein is PSPC1-SFPQ hetero-dimer. Lane 1: no PSPC1-SFPQ; lane 2: 500 nM PSPC1-SFPQ; lane 3: 1.5 μM PSPC1-SFPQ; lane 4: 4.5 μM PSPC1-SFPQ; lane 5: 13 μM PSPC1-SFPQ. E The DNA used is C10ss and the tested protein is PSPC1-SFPQ hetero-dimer. Lane 1: no PSPC1-SFPQ; lane 2: 500 nM PSPC1-SFPQ; lane 3: 1.5 μM PSPC1-SFPQ; lane 4: 4.5 μM PSPC1-SFPQ; lane 5: 13 μM PSPC1-SFPQ.

To characterize the PSPC1-HC and (PSCP1-SFPQ)-HC nucleoprotein complexes, we purified truncated forms of PSPC1 and SPFQ proteins known to assemble into dimers [24,25]: the truncated PSCP1 homo-dimer was from amino acids 53 to 320 and the truncated SFPQ protein in the PSCP1-SFPQ hetero-dimer was from residue 276 to 535 (indicated as double arrows underneath each schematic representation of the protein on Fig 3A). The choice to purify truncated forms of the proteins stems from the difficulty to purify the full length proteins. Both forms of truncated dimers contained the two tandem RNA recognition motifs, the NonA/paraspeckle domain but had a truncated carboxy terminal coiled-coiled domain. The purified proteins were tested for their efficiency of binding to HC and to the circular single stranded DNA coming from the nicking of dsMC10 mini-circle, C10ss (Fig EV1). High concentrations of PSPC1 and PSCP1-SFPQ (> 1 μM) were required to make possible the binding of the proteins to the HC (Figs 3B and 3D) whereas within these ranges of concentration, binding to C10ss was highly efficient (Figs 3C and 3E). For instance, at 9 μM PSPC1, 37% +/− 1.5% of the HC was engaged in a complex with PSPC1 (Lane 5 of Fig 3B) whereas the extent of assembly of PSPC1 on C10ss reached 95% +/− 1.5 % (Lane 5 of Fig 3C). These results indicated that the binding of the truncated forms of PSPC1 homo-dimer and PSCP1-SFPQ heterodimer to HC was not highly specific. It is possible that the full length forms of the homo-and hetero-dimers and/or their post-translational modifications mainly carried by the intrinsically disordered amino- and carboxy-terminal extremities make them acquire specificity for HC structures. Alternatively, proteins present in the A10 or A11 gel filtration fractions may be required to confer HC specificity to PSPC1 homo-dimer and PSCP1-SFPQ hetero-dimer.

### Detection of the SND1-HC complex on a sucrose gradient

SND1 belongs to the group of proteins that was partially purified from the Hela cell nuclear extracts as a HC-specific binding protein (Table 1). To confirm this interaction that was detected by a electrophoresis on a polyacrylamide under native conditions, we investigated whether the SND1-HC complex could also be isolated on a sucrose gradient. A 5 to 40% sucrose gradient of 5 mL was poured and the mixture of the interaction between radiolabeled HC and the fraction A10 of the size exclusion chromatography was loaded on the gradient. After centrifugation, fractions of 200 μL were collected from the top to the bottom of the gradient and analyzed for their DNA and SND1 content. DNA content was estimated by radioactivity measurement and SND1 content by western blot. Results indicate that SND1 peaked at fraction 8 when the fraction A10 of the size exclusion chromatography was sedimented alone (white bars on the graph of Fig 2D) where as its peak was shifted to fraction 5 when the fraction A10 was sedimented with HC (hatched bars on the graph of Fig 2D). Analysis of the radioactivity in fractions 5 to 10 of the sucrose gradient indicated that sedimentation profile of HC was also modified by the proteins in the fraction A10 and that fraction 5 did contain radiolabeled HC (Fig 2E). Altogether, the sucrose gradient sedimentation supported the formation of a complex between SND1 and HC.

### Characterization of the SND1-HC nucleoprotein complex with purified proteins

SND1 has multiple protein- and nucleic acid- binding partners (for reviews see [26–28] and is involved in many aspects of gene expression (*e.g.* transcription [29–35], RNA splicing [36,37], RNA interference [38,39], microRNA processing and decay [40,41], RNA protection in stress granules [42,43], protein modification and degradation [44,45]. While fulfilling these functions, it may have either a scaffolding role or a catalytic (nuclease) activity. Its domain composition comprises at the amino terminus of the protein, four staphylococcal nuclease (SN)-like domains assembled in tandem (SN1 to SN4 on Fig 5A) and at the carboxy terminal end of the protein a fusion of a Tudor domain with a partial SN domain (SN5 on Fig 4A).

**Figure 4.**
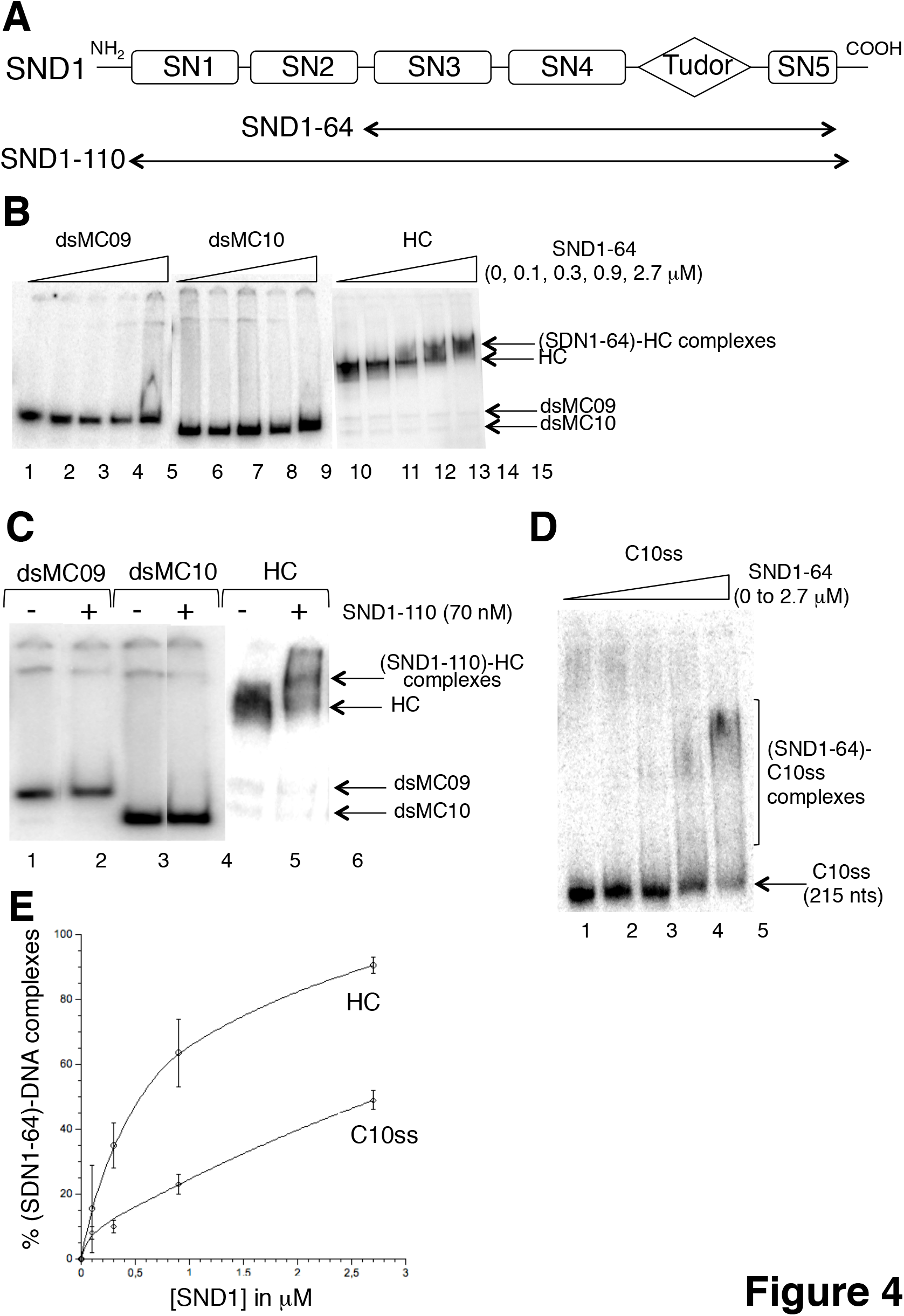
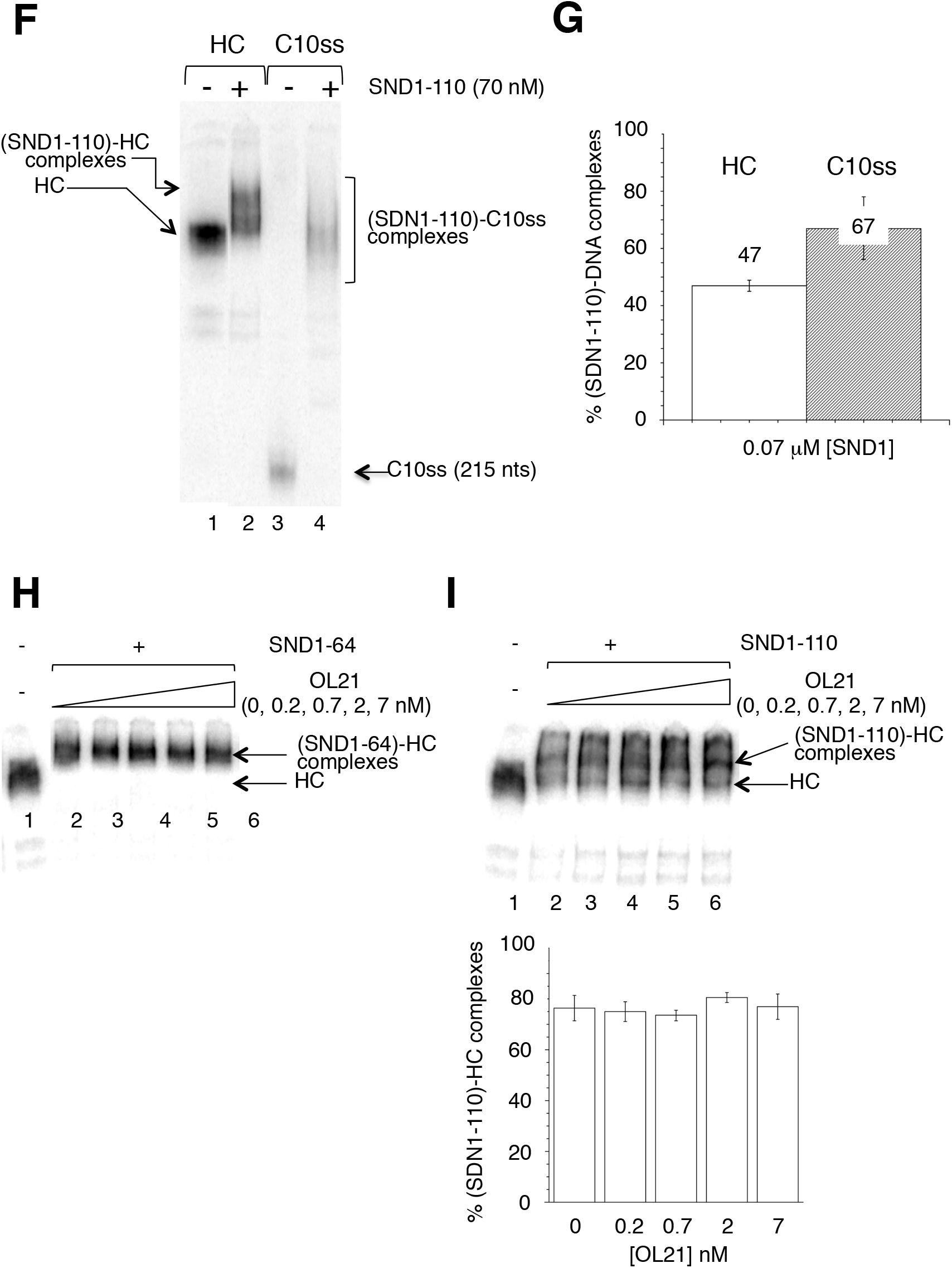
Interaction between purified SND1 proteins and various DNA constructs. A The organization of SND1 in domains is shown. The double arrow underneath the schematic representation corresponds to the proteins that were expressed in *E. coli* and purified. The name of the purified protein is indicated on the left side of the double arrow. B Interactions were performed in a final volume of 7.5 μL with 0.1 femtomole of radiolabeled DNA and the indicated amount of purified protein. Species were resolved by electrophoresis under native conditions. Three DNAs were tested for their binding to SND1-64: dsMC09 (lanes 1-5); dsMC10 (lane 6-10); HC (lanes 11-15). Concentrations of protein were as indicated (lanes 1, 6, 11: 0; lanes 2, 7, 12: 0.1 μM; lanes 3, 8, 13: 0.3 μM; lanes 4, 9, 14: 0.9 μM; lanes 5, 10, 15: 2.7 μM). Free DNAs and bound DNAs are indicated. C Interactions were performed in a final volume of 13.25 μL with 0.1 femtomole of radiolabeled DNA and the SDN1-110 at 70 nM. Species were resolved by electrophoresis under native conditions. Three DNAs were tested for their binding to SND1-110: dsMC09 (lanes 1 and 2); dsMC10 (lanes 3 and 4); HC (lanes 5 and 6). Free DNAs and bound DNAs are indicated. D Interactions were as described in (A). The DNA was C10ss, the single stranded circle obtained after nicking of the dsMC10 with Nt.BbvcI. 0.1 femtomole of C10ss was included in the reaction mixture and the concentrations of protein were as indicated (lane 1: 0; lane 2: 0.1 μM; lane 3: 0.3 μM; lane 4: 0.9 μM; lane 5: 2.7 μM). Free DNAs and bound DNAs are indicated. E The two curves (one for HC and one for C10ss) show the percentage of (SDN1-64)-DNA complexes as a function of protein concentration. Error bars correspond to the standard deviation. Percentages are the mean of three independent experiments. F Interactions were as described in (C). C10ss is the single stranded circle obtained after nicking of the dsMC10 with Nt.BbvcI. 0.1 femtomole of C10ss or HC was included in the reaction mixture and the concentration of protein was 70 nM. Free DNAs and bound DNAs are indicated. G The plot shows the percentage of (SDN1-110)-DNA complexes assembled at 70 nM SND1-110. Error bars correspond to the standard deviation. Percentages are the mean of three independent experiments. H The interaction between radiolabeled HC and SND1-64 was tested in the presence of OL21, an oligonucleotide long of 21 nucleotides. SDN1-64 was at 2.7 μM and HC at 14 pM. HC was premixed with increasing amount of OL21 (lane 1: no OL21, no SND1-64; lane 2: no OL21; lane 3: 0.2 nM OL21; lane 4: 0.7 nM OL21; lane 5: 2 nM OL21; lane 6: 7 nM OL21) before adding SND1-64. Free DNAs and bound DNAs are indicated. I The interaction between radiolabeled HC and SND1-110 was tested in the presence of OL21, an oligonucleotide long of 21 nucleotides. SDN1-110 was at 70 nM and HC at 7.5 pM. HC was premixed with increasing amount of OL21 (lane 1: no OL21, no SND1-110; lane 2: no OL21; lane 3: 0.2 nM OL21; lane 4: 0.7 nM OL21; lane 5: 2 nM OL21; lane 6: 7 nM OL21) before adding SND1-110. Species are separated by electrophoresis on a polyacrylamide gel. Free DNAs and bound DNAs are indicated. The plot shows the percentage of (SND1-110)-HC complexes as function of concentration of OL21. The standard deviation is calculated from two independent experiments.

**Figure 5.**
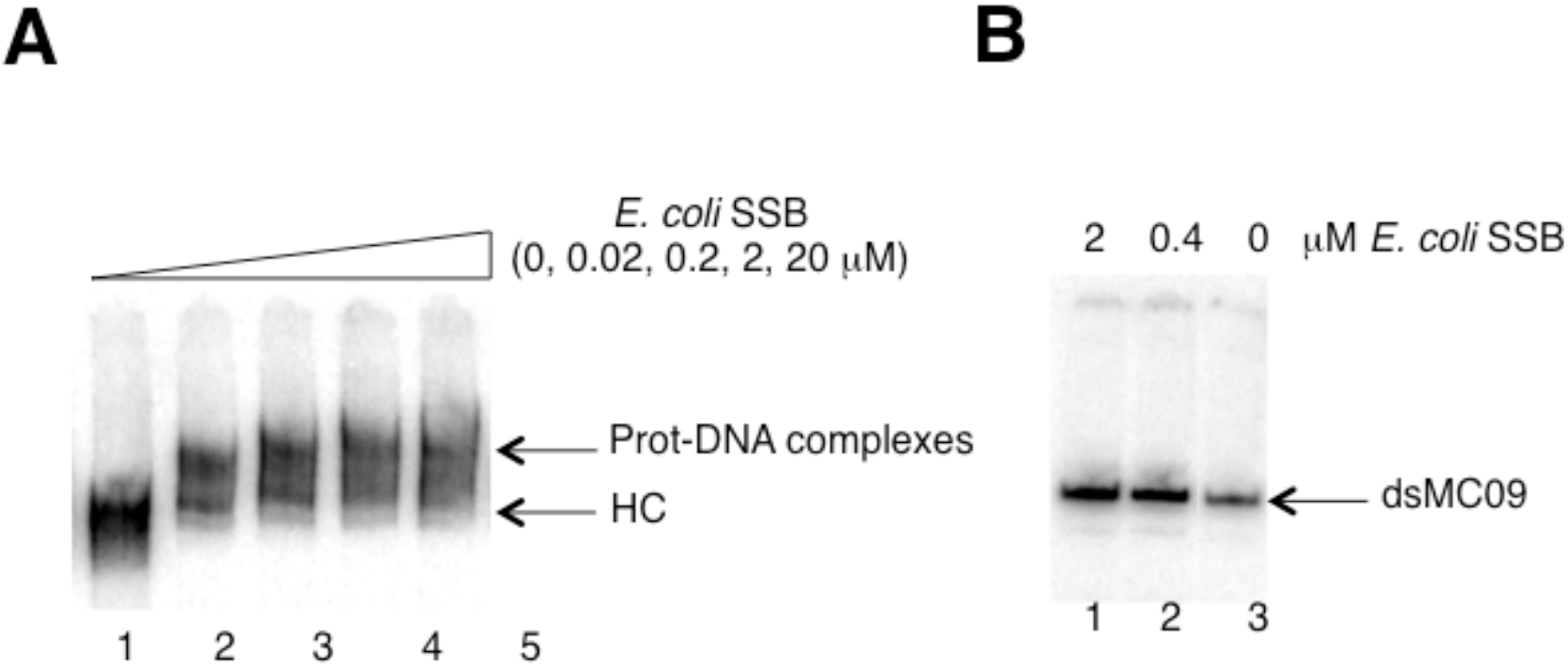
Interaction between *E. coli SSB* and DNA. Interactions were performed in a final volume of 7.75 μL with 0.1 femtomole of radiolabeled DNA and the indicated amount of purified protein. Species were resolved by electrophoresis under native conditions. Free DNAs and bound DNAs are indicated. A The DNA used was HC. Lane 1: no *E. coli* SSB; lane 2: 20 nM *E. coli* SSB; lane 3: 200 nM *E. coli* SSB; lane 4: 2 μM *E. coli* SSB; lane 5: 20 μM *E. coli* SSB. B The DNA used is dsMC09. Lane 1: 2 μM *E. coli* SSB; lane 2: 0.4 μM *E. coli* SSB; Lane 3: no *E. coli* SSB

We purified two versions of this protein. SND1-64, from amino acids 315 to 863, lacks the first two SN domains and the 20 last amino acids and SND1-110, from amino acids 33 to 888, is truncated in the first SN domain since it lacks the 32 first amino acids (Fig 4A). We first characterized the interaction between SND1-64 and two types of DNA constructs (Fig 4B): the HC and the dsMCs that were used to prepare the HC, dsMC09 and dsMC10. Results indicated that over the range of concentration tested (from 0 to 2.7 μM) SND1-64 did interact with HC (lanes 11 to 15 of Fig 4B) but not with dsMC (lanes 1 to 10 of Fig 4B). The same result was obtained with SDN1-110 although a single concentration of protein (70 nM) was tested due to the low concentration of SDN1-110 that was purified (Fig 4C). Quantification of the gels indicated that SND1-110 exhibited a stronger affinity to HC than SND1-64 (compare Figs 4E and 4G), possibly due to the presence of two additional SN domains at the amino terminal side of SND1-110.

Single stranded DNA might be exposed at the junction of the HC strands and be the substrate recognized by SND1 proteins. To check this hypothesis, we first characterized the interaction between HC and *E. coli* SSB, the single stranded DNA binding protein from *E. coli* (Fig 5). Results showed that *E. coli* SSB interacted with the HC (Fig 5A) but not with dsMC09 (Fig 5B), indicating that the HC molecule carried regions of single stranded DNA. As it was possible that SDN1 only recognized the single stranded DNA of the HC and not the junctions of the strands of the HC, we performed two types of additional experiments. We first compared the binding of SND1 to HC and the binding of SND1 to C10ss (the circular single stranded DNA coming from the nicking of dsMC10, Fig EV1). Results showed that SND1-64 did bind C10ss but the binding was weaker than that measured with HC (Figs 4D-E). In the second type of experiment, we investigated whether a single stranded oligonucleotide long of 21 nucleotides (OL21) could compete with HC for the binding of SND1. To that end, we mixed the HC substrate (10 pM) with increasing amount of OL21 (from 0 to 7 nM), added SND1-64 to the mixture and, after 25 minutes of incubation at 22°C, resolved the species by electrophoresis on a polyacrylamide gel under native conditions. Results indicated that the (SND1-64)-HC complex was highly resistant to the presence of the OL21 competitor since all the HC could still be bound by SND1-64 even with a 700 fold molar excess of competitor (Fig 4H). This competition experiment confirmed that the single stranded DNA was not the only binding substrate recognized by SND1-64 on the HC. We performed the same kind of experiments with the SND1-110 protein. Results indicated that SND1-110 bound almost equally well HC and C10ss DNAs (Figs 4F-G) and that OL21 oligonucleotide poorly competed with HC to bind to SND1-110 (Fig 4I).

## Conclusion

In this work we identified SND1 as a protein exhibiting a high specificity for the four strand junction of the HC. It is known that SND1 protein interacts with various proteins of the transcription machinery, including the RNA polymerase II [35] and the transcription initiation factor TFIIE [34]. Therefore, it is tempting to propose that the binding of SND1 to HCs participates into some essential functions of SND1, including the function of regulation of gene expression. Considering the proposed role of HCs as potential structures affecting dynamically genome organization and transcription [19], SND1 could for example act by bridging hemicatenated structures and the basal transcription machinery. The dynamics of HC structures, whose location may change with time, cell types or upon external stresses, could therefore be modulated by specific binding of SND1. Given the essential role of the SND1 in transcription regulation and in oncogenic progression, our results invite to explore further the interaction between SND1 and HCs.

## Materials and Methods

### Materials

The Hela nuclei were from Ipracell and were resuspended in 10 mM Hepes pH 7.9, 1.5 mM MgCl_2_, 10 mM KCl, 0.5 mM DTT. Oligonucleotides were from Eurogentec. T4 polynucleotide kinase, T4 DNA ligase, Nt.BbvcI and Nb.BbvcI nicking enzymes were from New England Biolabs. Wheat germ topoisomerase was from Inspiralis. *E. coli* SSB was from Ubs. γ^32^P-ATP was from Perkin Elmer. Trypsin was from Promega (USA). Cellulose phosphate was from Sigma. The HiTrap ButylFF column of 1 mL and the gel filtration column Superdex 200 10/300 GL were from GE Healthcare. MacroPrep HighS resin was from Biorad. All columns were prepared as recommended by the manufacturer and equilibrated in the indicated loading buffer before use. λ DNA was from Sigma. Acrylamide and bisacrylamide stock solutions were from Euromedex. Proteinase inhibitor cocktail was from Roche. Anti-SND1 and anti-PTPB1 antibodies were from St John’s Laboratory. Anti-SFPQ antibody was from Novusbio. Anti-PSPC1 was from Abcam. The sequence of the oligonucleotide OL21 (from Eurogenetec) was: 5’-CCCTAACCCTAACCCTAACCC-3’.

### Preparation of HCs

HCs were prepared as described in [20] with three modifications in the procedure. Firstly, the production of dsMCs was performed in the presence of ethydium bromide (0.35 μg/mL) to favor circularization of DNA fragments into monomeric circles. Secondly, dsMCs were purified on a 4% acrylamide (29:1 = acrylamide:bisacrylamide mass ratio) gel made in TBE 0.5X before performing the nicking reaction. Thirdly, after the nicking reaction, the circular and linear single stranded fragments were purified on a 4% acrylamide (29:1 = acrylamide:bisacrylamide mass ratio) gel made in TBE 0.5X + 7M Urea.

### Preparation of protein extract from Hela nuclei

6 mL of lysis buffer containing 10 mM Hepes pH 7.8, 0.6 M NaCl, 1.5 mM MgCl_2_, 5 mM DTT, 0.8 mg/mL PMSF and a cocktail of proteinase inhibitors were added to the 12 mL containing 15×10^9^ Hela nuclei and left on ice until thawed. While gently vortexing the nuclei, EDTA was added to a final concentration of 2.5 mM and 410 μL of NaCl 5 M were added four times at intervals of five minutes. The 0.6 M NaCl final concentration that was reached at the end permitted to the chromatin to gently swell. The sample was left on ice for 45 minutes and centrifuged 60 minutes at 4°C and at 184000 g. The supernatant of the centrifugation that constituted the nuclei protein extract was next fractionated by different means (see “Fractionation of nuclei proteins” section). Each step of fractionation generated fractions that were tested for containing HC-specific binding proteins (see “Test of chromatographic fractions for containing HC-specific binding activity”). Interesting fractions were next combined and applied to the next step of the fractionation procedure.

### Test of chromatographic fractions for containing a HC-specific binding activity

Unless indicated otherwise, the test consisted of performing an interaction between DNA (HC or dsMC) and chromatography fractions by mixing 8 μL of each fraction with 1 μL of DNA (0.1 femtomole) in a 1.5 mL Eppendorf tube. Competitor λ DNA (48502 base pairs) was added at the indicated concentration (from 0 to 30 ng (1 femtomole) of DNA molecule). The mixtures were incubated for 30 minutes at 22°C. At the end of the incubation glycerol was added to a final concentration of 5%. The samples were loaded on a 4% acrylamide (29:1 = acrylamide:bisacrylamide mass ratio) gel made in Tris-Borate 89 mM, boric acid 89 mM EDTA 2 mM (TBE) 0.5X. Electrophoresis was performed at 4°C for 4 hours and at 150 V. The gel was dried and exposed on a ^32^P sensitive screen. The fractions that permitted the HC (and not the dsMC) to shift by decreasing its mobility in the gel were considered as containing HC-specific binding proteins.

### Fractionation of nuclei protein extract

The first step of protein fractionation was a protein precipitation that was performed with 5% of ammonium sulfate. Ammonium sulfate was slowly added to the protein extract under slow stirring and at 4°C. Proteins were let precipitated overnight. Precipitated proteins were recovered by centrifugation (20 minutes, 4°C, 1200 g). The pellet of proteins was resolubilized in 27 mL of a low salt (Low KP) buffer containing 25 mM potassium phosphate pH 7.5, 2 mM EDTA, 10% glycerol, 12.5 mM DTT. The resolubilized proteins were next loaded at a 0.5 mL/min flow rate on a column of cellulose phosphate of 4 mL equilibrated in the Low KP buffer. After the load of proteins, the column was washed with 20 mL of Low KP buffer. The bound proteins were eluted with a high salt buffer containing 500 mM potassium phosphate pH 7.5, 2 mM EDTA, 10% glycerol, 12.5 mM DTT and contained HC-specific binding proteins. They were combined and dialyzed against a high salt buffer (High KP) containing 500 mM potassium phosphate pH 7.5, 2 mM EDTA, 10% glycerol, 2 mM DTT. After dialysis, the protein sample was loaded at a flow rate of 0.3 mL/min on the 1 mL HighTrap Butyl-FF column equilibrated with the High KP buffer. The flow through of the column was collected and dialyzed overnight against a low salt buffer containing 25 mM potassium phosphate pH 6.6, 25 mM KCl, 2 mM EDTA, 10% glycerol, 2 mM DTT. After dialysis, the protein sample was loaded on a 1 mL MacroPrep High S at a flow rate of 0.7 mL/min. After loading, the column was washed with 15 mL of the low salt buffer (25 mM potassium phosphate pH 6.6, 25 mM KCl, 2 mM EDTA, 10% glycerol, 2 mM DTT). The proteins were eluted with a 15 mL salt gradient (KCl raising from 25 mM to 500 mM). Fractions containing HC-specific binding proteins were pooled, concentrated and loaded on a Superdex 200 (10/300 GL) equilibrated in a buffer containing 25 mM potassium phosphate pH 7.5, 150 mM KCl, 2 mM EDTA, 10% glycerol, 2 mM DTT. Fractions of interest were kept at −80°C in small aliquots.

### Sample preparation for mass spectrometry analysis

A large scale interaction between 16 femtomoles of HC and the designated chromatography fraction was performed in a final volume of 150 μL of buffer containing 7.5 mM potassium phosphate buffer pH 7.5, 34 mM Tris HCl pH 7.5, 45 mM KCl, 34 mM NaCl, 1 mM EDTA, 0.03 % Triton X100, 3% glycerol, 2 mM DTT. When present competitor λ DNA was added at the indicated concentration. The control sample that was included in the analysis consisted of the protein fraction alone (no HC) made in the same interaction buffer. The mixtures were next incubated for 30 minutes at 22°C. At the end of the incubation glycerol was added to a final concentration of 5%. The samples were loaded on a 4% acrylamide (29:1 = acrylamide:bisacrylamide mass ratio) gel made in TBE 0.5X. Electrophoresis was performed at 4°C for 4 hours and at 150 V. After electrophoresis the gel was exposed on a ^32^P sensitive screen. The material (as gel cubes manually excised from the gel) that migrated above the HC was analyzed by mass spectrometry for its protein content.

### Protein identification by LC-MS/MS analysis

In-gel digestion was carried out with trypsin: sample were washed twice with a mixture of 100 mM ammonium bicarbonate (ABC) and 50% (vol/vol) acetonitrile (ACN) for 20 min at room temperature and then dehydrated using 100% ACN for 20 min, before being reduced with 25 mM ABC containing 10 mM DTT for 1 h at 56 °C and alkylated with 55 mM iodoacetamide in 25 mM ABC for 30 min in the dark at room temperature. Gel pieces were washed twice with 25 mM ABC and dehydrated (twice, 20 min) with 100% ACN. Gel cubes were incubated with sequencing grade-modified trypsin (12.5 ng/μl in 40 mM ABC with 10% ACN, pH 8.0) overnight at 37 °C. After digestion, peptides were extracted twice from gel pieces with a mixture of 50% ACN – 5% formic acid (FA) and then with 100% ACN. Extracts were dried using a vacuum centrifuge concentrator plus (Eppendorf).

Mass spectrometry (MS) analyses were performed on a Dionex U3000 RSLC nano-LC system coupled to an Orbitrap Fusion Tribrid mass spectrometer (Thermo Fisher Scientific). After drying, peptides were solubilized in 7 μL of 0.1 % trifluoroacetic acid (TFA) containing 10 % acetonitrile (ACN). One μL was loaded, concentrated and washed for 3 min on a C_18_ reverse phase precolumn (3 μm particle size, 100 Å pore size, 75 μm inner diameter, 2 cm length, Thermo Fisher Scientific). Peptides were separated on a C_18_ reverse phase resin (2 μm particle size, 100 Å pore size, 75 μm inner diameter, 25 cm length from Thermo Fisher Scientific) with a 30 minutes gradient starting from 99 % of solvent A containing 0.1 % FA in H_2_O and ending in 90 % of solvent B containing 80 % ACN, 0.085 % FA in H_2_O. The mass spectrometer acquired data throughout the elution process and operated in a data-dependent scheme with full MS scans acquired with the Orbitrap, followed by MS/MS HCD fragmentations acquired with the Ion Trap on the most abundant ions detected in top speed mode for 3 seconds. Resolution was set to 60,000 for full scans at AGC target 2.0e5 within 60 ms maximum injection ion time (MIIT). The MS scans spanned from 350 to 1500 m/z. Precursor selection window was set at 1.6 m/z, and MS/MS scan resolution was set with AGC target 2.0e4 within 100 ms MIIT. HCD Collision Energy was set at 30 %. Dynamic exclusion was set to 30 s duration. For the spectral processing, the software used to generate .mgf files was Proteome Discoverer 1.4 (ThermoFisher Scientific). The mass spectrometry data were analyzed using Mascot v2.5 (Matrix science) on *Homo sapiens* (20,243 sequences) from the SwissProt databank containing 553,655 sequences; 198,177,566 residues (April 2017). The enzyme specificity was Trypsin’s and up to 1 missed cleavage was tolerated. The precursor mass tolerance was set to 4 ppm and the fragment mass tolerance to 0.55 Da for Fusion data. Carbamidomethylation of cysteins and oxidation of methionines were set as variable modifications. Among positive identifications based on a Mascot score above the significance threshold p<5%, we selected proteins identified with at least 2 peptides with an ion score > 25 for each of them.

### Sucrose gradient

The solution used to prepare the sucrose gradient contained 50 mM Tris HCl p7.5, 50 mM NaCl, 0.5 mM EDTA, 2 mM DTT and either 5 % or 40 % sucrose. A 5 mL gradient of 5 to 40 % sucrose was prepared with a gradient maker and a peristaltic pump and equilibrated at 8°C for two hours before use. The interaction between HC and the designated fraction from the gel filtration was performed in a 90 μL volume and its product was carefully loaded on top of the sucrose gradient. Two control samples were also run at the same time: a sample containing only the HC and a sample containing only the designated fraction from the gel filtration. The centrifugation was performed at 8°C, 64090g in SW 55Ti rotor for 12 hours. At the end of the run, fractions of 200 μL were recovered from the top to the bottom of the gradient and tested for the presence of (i) HC by measuring their reactivity level and (ii) the SND1 protein by western blot.

### Western Blot

Western blot was performed with nitrocellulose membrane and in TBS buffer supplemented with 0.1 % Tween20 and 5 % low fat milk. Anti-PSPC1 and anti-SND1 antibodies were used at a final concentration of 1 μg/mL, anti-SFPQ at 0.4 μg/mL (final concentration) and anti-PTBP1 at 0.1 μg/L (final concentration).

### Purification of SND1 proteins

Plasmids overexpressing the human Tudor-Staphylococcal Nuclease-like protein from amino acids 33 to 888 (SND1-110) and from amino acids 315 to 863 (SND1-64) were a gift from Pr. Hanna S. Yuan. The purification of the two proteins was performed as described in [38].

### Purification of PSPC1 homo-dimer and PSPC1-SFPQ hetero-dimer

Plasmids overexpressing PSPC1 protein from residues 53 to 320 and SFPQ from residues 276 to 535 were a gift from Pr. Mihwa Lee. The purification of PSPC1 was performed as described in the thesis manuscript of Daniel Michael Passon [25]. The purification of the PSPC1-SFPQ heterodimer was performed as described in [24].

### Interaction between DNA and purified proteins

Unless mentioned otherwise, the interaction between 0.1 femtomole of DNA (radiolabeled HC, radiolabeled dsMC or radiolabeled circular single stranded DNA C10ss) and the purified proteins was performed in 8 μL of TENT buffer (10 mM Tris HCl pH pH 7.5, 50 mM NaCl, 0.5 mM EDTA, 0.05 % Triton X100) supplemented with 2 mM DTT. The concentrations of proteins were as indicated in the figure legend. The mixtures were incubated for 30 minutes at 22°C. At the end of the incubation glycerol was added to a final concentration of 5 %. The samples were loaded on a 4 % acrylamide (29:1 = acrylamide:bisacrylamide mass ratio) gel made in TBE 0.5X. Electrophoresis was performed at 4°C for 3 hours and at 150 V. The gel was dried and exposed on a ^32^P sensitive screen. After exposure, the screen was scanned and quantification of the gel was performed using the ImageQuant TL Software.

## Acknowledgments

The research was supported by core funding of the laboratory by INSERM, CNRS, and the National Museum of Natural History. The authors thank members of the Genome Structure and Instability laboratory for helpful discussions. ED is particularly grateful to Dr François Strauss for introducing her to the field of hemicatenanes and for the support and constructive discussions along the course of the project.

## Author contributions

ED conceived the experiments, performed experiments except the sucrose gradient and western blotting experiments that were performed by OR. CB and FG performed the identification of the HC-binding proteins by LC-MS/MS analysis at the proteomic platform 3P5 (Université de Paris, Institut Cochin, INSERM, U1016, CNRS, UMR8104, F-75014 Paris, France). All authors provided detailed and fruitful comments.

## Conflict of interest

The authors declare that they have no conflict of interest.

## Expanded View Figure Legend

### Figure EV1. Scheme of construction of DNA HCs from two dsMCs

Radiolabeled dsMCs09 and dsMCs10 were prepared from ^32^P -end labeled linear double stranded fragments of 215 (colored in red) and 235 (colored in blue) base pairs. Their nicking by specific enzymes led to two linear single stranded fragments (L09ss in blue and L10ss in red) and two single stranded circles (C09ss in blue and C10ss in red). Single stranded catenanes were assembled from the circularization of L09ss and L10ss under specific conditions: the linker oligonucleotide complementary to 25 nucleotides of L09ss and 25 nucleotides of L10ss brings together the two strands and favors the catenation. The single stranded catenanes were gel-purified and re-annealed with the C09ss and C10ss using the wheat germ Topoisomerase I to finally get the DNA HCs. Adapted from [20].

